# Inferring the probability of the derived versus the ancestral allelic state at a polymorphic site

**DOI:** 10.1101/257246

**Authors:** Peter D. Keightley, Benjamin C. Jackson

## Abstract

It is known that the allele ancestral to the variation at a polymorphic nucleotide site cannot be assigned with certainty, and that the most frequently used method to assign the ancestral state – maximum parsimony – is prone to mis-inference. Estimates of counts of sites that have a certain number of copies of the derived allele (the unfolded site frequency spectrum, uSFS) made by parsimony are therefore also biased. We previously developed a maximum likelihood method to estimate the uSFS for a focal species, using information from two outgroups and assuming simple models of nucleotide substitution. Here, we extend this approach to infer the uSFS, allowing multiple outgroups, potentially any phylogenetic tree topology and more complex models of nucleotide substitution. We find, however, that two outgroups and assuming the Kimura 2-parameter model is adequate for uSFS inference in most cases. We show that using parsimony for ancestral state inference at a specific site seriously breaks down in two situations. The first is where the outgroups provide no information about the ancestral state of variation in the focal species. In this case, nucleotide variation will be under-estimated if such sites are removed from the data. The second is where the minor allele in the focal species agrees with the allelic state of the outgroups. In this situation, parsimony tends to over-estimate the probability of the major allele being derived, because it fails to account for the fact that sites with a high frequency of the derived allele tend to be rare in most data sets. We present a method that corrects this deficiency, which is capable of providing unbiased estimates of ancestral state probabilities on a site-by-site basis and the uSFS.

## Introduction

Many population genetic and quantitative genetic analysis methods require the assignment of ancestral *versus* derived states at polymorphic nucleotide sites. For example, Fay and Wu (2000) and Zeng et al. (2006) proposed statistics, *H* and *E*, that compare the numbers of high, intermediate and low frequency derived variants, and these can be used to distinguish between different modes of natural selection and demographic change. A number of methods have also been developed to infer selection and demographic change based on the complete distribution of counts of derived alleles across sites, the unfolded site frequency spectrum (uSFS) (e.g., Boyko et al. 2008; Schneider et al. 2011; Tataru et al. 2017).

The minor allele at a site or counts of numbers of minor alleles at a group of sites (the folded site frequency spectrum) can be observed directly from sequence polymorphism data. In contrast, the derived *versus* the ancestral allele at a site cannot be known with certainty, because at least one outgroup is required for inference, and there is the possibility of more than one mutation separating the focal species from the outgroup. This implies that the uSFS also cannot be known precisely. For the purpose of ancestral state inference, rule-based maximum parsimony using outgroup species is the most frequently applied method in molecular evolutionary genetics (e.g., Voight et al. 2006; Dreszer et al. 2007; Keinan et al. 2007; Sabeti et al. 2007; Lohse et al. 2011; Langley et. 2012; 1000 Genomes Project Consortium 2010, 2015; Schmidt et al. 2017). It has been recognized, however, that parsimony potentially produces misleading results (Felsensein 1981; Eyre-Walker 1998; Collins et al. 1994). Of particular relevance here is that sites that have a low frequency of the derived allele are usually more common than sites that have a high frequency of the derived allele. This implies that misinference tends to lead to upwardly biased counts of high frequency derived alleles (Baudry and Depaulis 2003; Hernandez et al. 2007).

There is a related problem concerning the assignment of ancestral states at polymorphic sites, which does not seem to have been addressed. If ancestral states are assigned site-by-site, potentially useful information is ignored. For example, in Fig. 1 the single outgroup species (state G) is uninformative about the ancestral allele of the variants in the focal species (which must be T or A). It is more likely, however, that the ancestral allele is T, if sites with a high frequency of the derived allele are uncommon in the dataset as a whole (as is usually the case).

**Figure 1.**
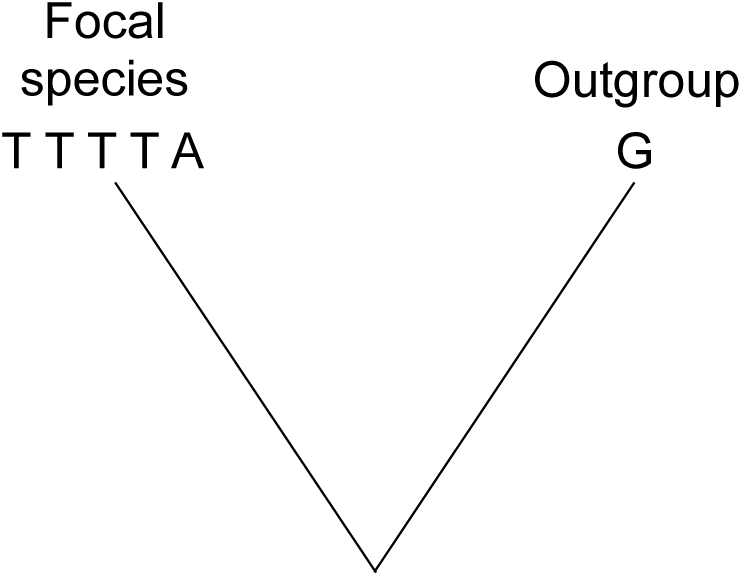
A site has *n* = 5 gene copies in a focal species. The major and minor alleles are T and A, respectively. The single outgroup (state G) is uninformative about the ancestral state of the variation in the focal species.

Matsumoto et al. (2015) pointed out that ancestral states are not observable, that a single best ancestral reconstruction is not advisable, and that assuming one can bias molecular evolutionary inference. This was developed by Jackson et al. (2017), who assigned the ancestral state probability at a focal site as the inferred probability of the node for the common ancestor of the focal species and the closest outgroup, obtained using PAML, Yang (2007), ignoring polymorphism data. This does not optimally weight information coming from the focal site itself and from the complete set of sites in the data.

Inference of ancestral states on a site-by-site basis has been problematical, but there has been progress in inferring the uSFS. Hernandez et al. (2007) developed a context-dependent substitution model using a single outgroup to infer the ancestral state at a polymorphic site in a focal species, then implemented a step to correct for ancestral misidentification. Their simulations suggested, however, that the approach only partially corrects for ancestral misidentification, depending on the divergence between the focal species and the outgroup.

Schneider et al. (2011) developed a probabilistic method to infer the uSFS on a site-by-site basis, but did not use information from the frequencies of polymorphisms across all sites, so results from this method are biased. Keightley et al. (2016) developed a maximum likelihood (ML) method (ml-est-sfs) that addresses the deficiency in Schneider et al. (2011), and simulations suggested that it is capable of correctly inferring the uSFS. It uses a two-stage process, in which the evolutionary rates are estimated by ML, then, assuming the rates, estimates the uSFS elements by ML, while correctly weighting information from informative and uninformative sites. However, the method is limited to two outgroups, assumes simple substitution models (for one outgroup, the Kimura 2-parameter model and for two outgroups, the Jukes-Cantor model), and is not readily scalable to more than two outgroups or more to complex substitution models. It is unknown whether more realistic substitution models and/or further outgroups significantly improves inference accuracy. Furthermore, it does not assign ancestral state probabilities on a site-by-site basis.

In this paper, we further develop the approach of Keightley et al. (2016), with the following objectives. 1. Estimate the uSFS, allowing several outgroups, potentially any tree topology, and realistic nucleotide substitution models. 2. Infer ancestral state probabilities for each polymorphic site in the data. We evaluate the performance of new approach by simulations, apply it to data from the Drosophila Population Genomics Project (DPGP) as a test case, and re-infer the ancestral state probabilities for the 1000 Genomes Project in humans, which were previously inferred by a parsimony-related approach.

## Materials and Methods

Following Keightley et al. (2016), uSFS inference is carried out in two-steps. Evolutionary rate parameters are estimated in step 1, then in step 2 the uSFS is computed conditional on the evolutionary rate parameter estimates. Information from steps 1 and 2 is then combined in a third step to infer the ancestral state probability for each polymorphic site.

### Representation of the data and some definitions

Suppose we have sampled *n* gene copies at a set of sites from a population of a focal species. The uSFS we require to estimate therefore contains *n* - 1 elements, excluding the elements where the ancestral or derived allele is fixed. We assume that a single gene copy at each site is known for one or more outgroup species, and the tree topology relating the species is known without error (Fig. 2). In the analysis we assume that the nucleotide variation within the focal species coalesces within the branch labeled b1. The consequences of violation of these assumptions is investigated in simulations. The observed nucleotide configuration for a site is the count of each of the four nucleotides in the focal species (labeled X, Y for a biallelic site), along with the state for each outgroup, represented by the presence of a single nucleotide A, C, G or T. Let the number of outgroups = *n*_*o*_ (in Fig. 1, *n*_*o*_ = 3), and denote the outgroups *o*_1_, *o*_2_…*o*_*n*_. Assuming an unrooted tree (as in Fig. 2), the number of branches in the tree is therefore *n*_*B*_ = 2*n*_*o*_ − 1.

**Figure 2.**
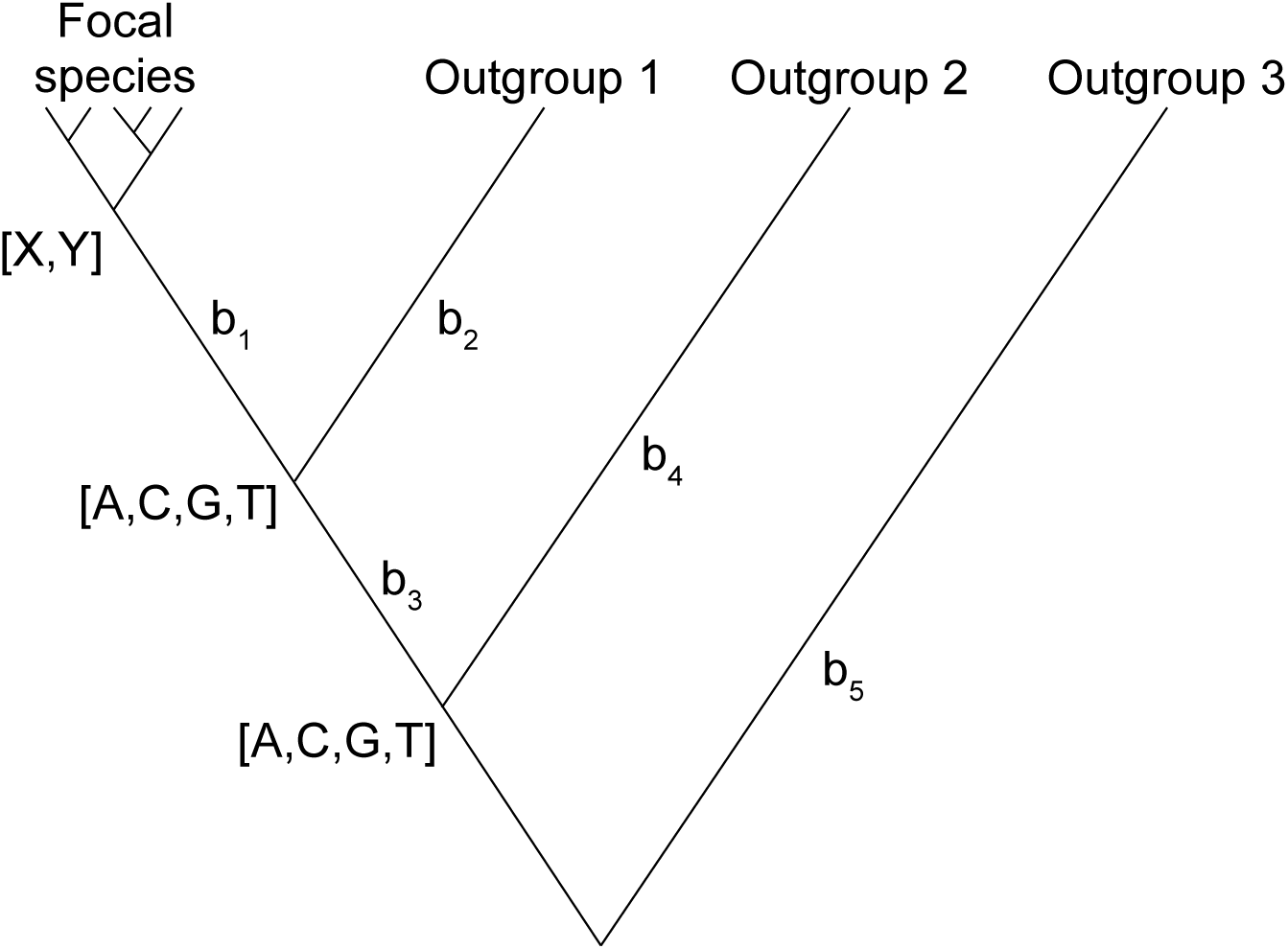
Representation of the data for uSFS and ancestral state inference. Polymorphism within the focal species (nucleotides X, Y) is assumed to coalesce within branch b_1_. There are three outgroups, two unknown internal nodes and five branches in this tree. The root of the tree is not identifiable, therefore branch b_5_ extends from outgroup 3 to the node of b_3_ and b_4_.

### Models of nucleotide substitution

The Jukes-Cantor model (JC), Kimura 2-parameter model (K2) and a model allowing six symmetrical rates (R6; Fig. 3a) are considered. All substitution models require the estimation of evolutionary rates (= mean number of nucleotide changes per site) for each branch, *K*_1_..*K*_*n*_*B*__. The rates are the only parameters for the JC model. For the K2 model, an additional parameter, *κ*, specifies the rate of transition mutations relative to the rate of transversions. For the R6 model there are six symmetrical relative mutation rates, *r*_1_..*r*_6_, 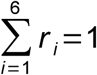 (Fig. 3a), so five independent parameters, *r*_1_..*r*_5_, require to be estimated.

**Figure 3.**
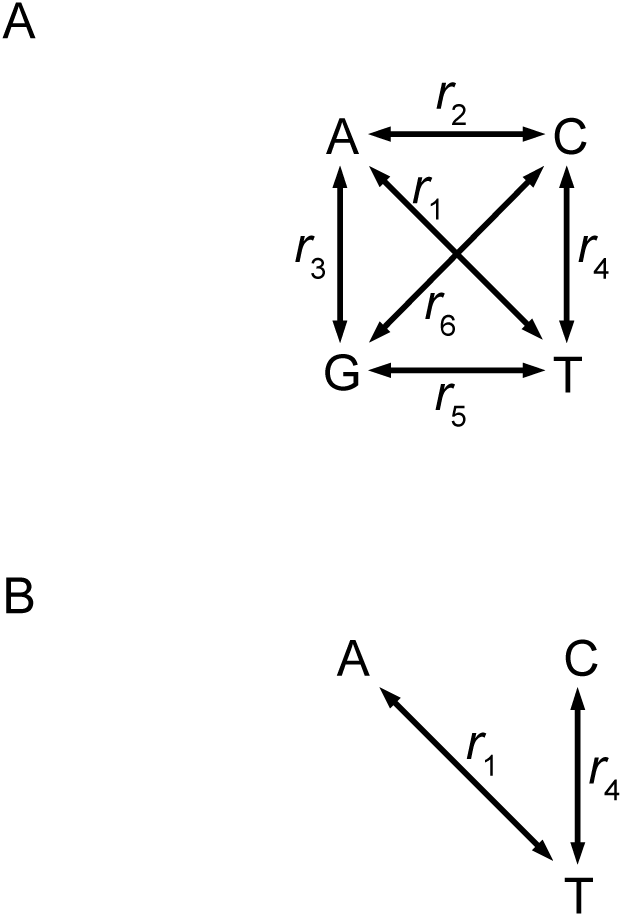
A The R6 model. B A simplified version of the R6 model used to illustrate the computation of probabilities of different numbers of changes on a branch, where all rates except *r*_1_ and *r*_4_ are zero.

### Estimation of rate parameters

Assuming the tree topology of Fig. 2, there are *n*_*B*_ substitution rates, and these, along with parameters of the substitution model (i.e., *κ* for the K2 model or *r*_1_..*r*_5_ for the R6 model), are estimated by maximum likelihood using the simplex algorithm for likelihood maximization (Nelder and Mead 1965). We checked convergence by picking starting values for the parameters from wide distributions, restarting the algorithm when convergence had apparently been achieved and checking that the same final maximum log likelihood was reached in multiple runs. Let **φ** be a vector specifying the model parameters, and let **y**_i_ be a vector specifying the observed nucleotide configuration for the focal species and the outgroups at site *i*. Sites are assumed to evolve independently, so the overall likelihood of the data is the product of probabilities of the observed nucleotide configuration for each site:

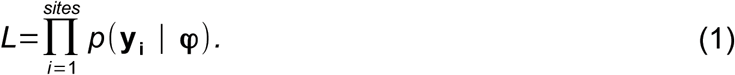

The probability of the nucleotide configuration for each site is evaluated by summing the probabilities for the *n*_*tree*_ = 4^*n*_*o*_-1^ possible unrooted trees, formed from all possible nucleotide combinations [A, T, G, C] at the unknown internal nodes along with the observed nucleotide configuration for the focal species and outgroups at the site (Fig. 2).

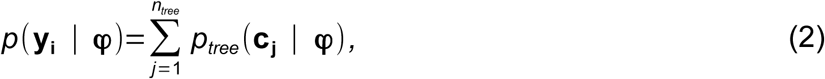

where *c*_*j*_is a vector representing the observed nucleotide configuration for the focal species and the *n*_*o*_ outgroups along with the nucleotide states for the *n*_*o*_ – 1 internal nodes for tree *j*. If the focal species is polymorphic at a site, the probability for that site is computed as the average probability for each observed nucleotide (X, Y in Fig. 2).

The overall probability for a given tree is computed from the product of the probabilities of each branch (*k* = 1..*n*_*B*_), conditional on the nucleotide states *x*_1, *k*_ and *x*_2, *k*_ representing the ancestral and derived nucleotides of that branch, given the nucleotide states specified in **c**_*j*_.

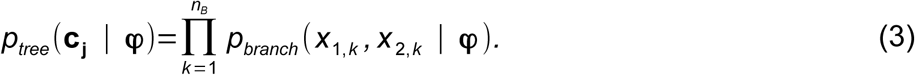

The probability for a branch depends on whether *x*_1, *k*_ and *x*_2, *k*_ differ from one another, the type of any difference (except in the case of the JC model), and the substitution rate parameters **φ**.

### Computation of *p*_*branch*_

In computing the probability of observing nucleotides *x*_1, *k*_ and *x*_2, *k*_ on branch *k*, it is assumed that the number of nucleotide changes on the branch is Poisson distributed. Terms for more than two changes on a branch are disregarded. Let *K*_*k*_ be the evolutionary rate parameter for branch *k*, which is the mean number of changes for that branch.

#### 1. JC model

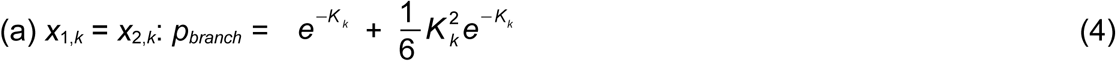

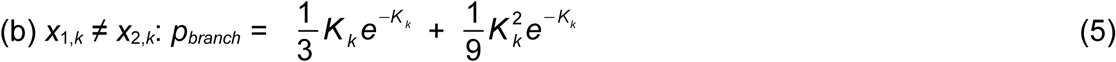

#### 2. K2 model

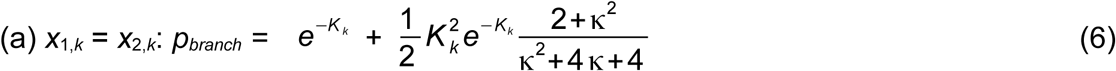

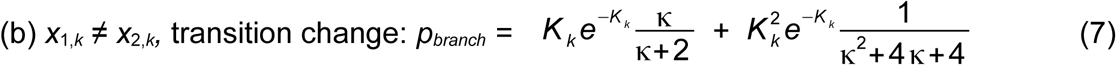

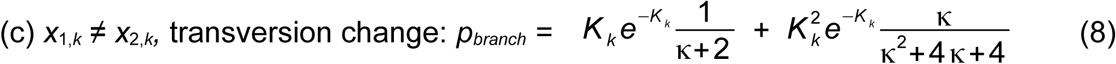

#### 3. R6 model (Fig. 3a)

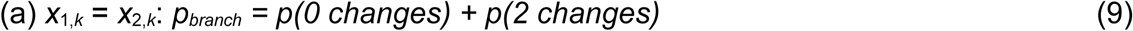

Taking the example of *x*_1, *k*_ = *x*_2, *k*_ = A:

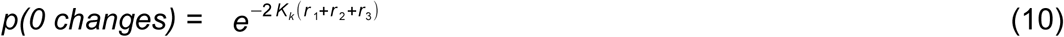

Note that *r*_1_, *r*_2_ and *r*_3_ are the relative rates for changes involving base A.

*p(2 changes)*:

The algorithm to compute the probability of observing the same ancestral and evolved base when two changes have occurred on a branch is illustrated by a simplified example where all relative rates in the model apart from two (*r*_1_ and *r*_4_) are zero (Fig. 3b).

For the case of *x*_1, *k*_ = *x*_2, *k*_ = A, the sequence of events must therefore be an A → T change followed by a T → A change. The probability of these events is obtained from:

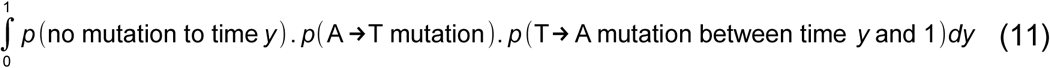

For the example illustrated in Fig. 3b, this is:

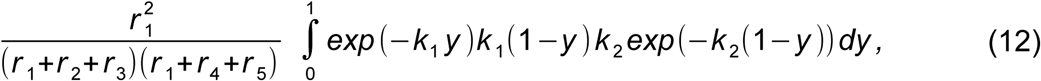

where *k*_1_ = 2*K*_*k*_(*r*_1_ + *r*_2_ + *r*_3_) and *k*_2_ = 2*K*_*k*_(*r*_1_ + *r*_4_ + *r*_5_). In this example the relative rates *r*_2_, *r*_3_ and *r*_5_ are all zero, but are included for completeness. Evaluation of the definite integral in (12) gives a closed form expression:

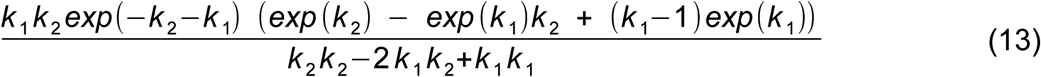

The logic can be extended to allow all the relative rates to be non-zero. (b) *x*_1, *k*_ ≠ *x*_2, *k*_: *p*_*branch*_ = *p*(1 change) + p(2 changes)

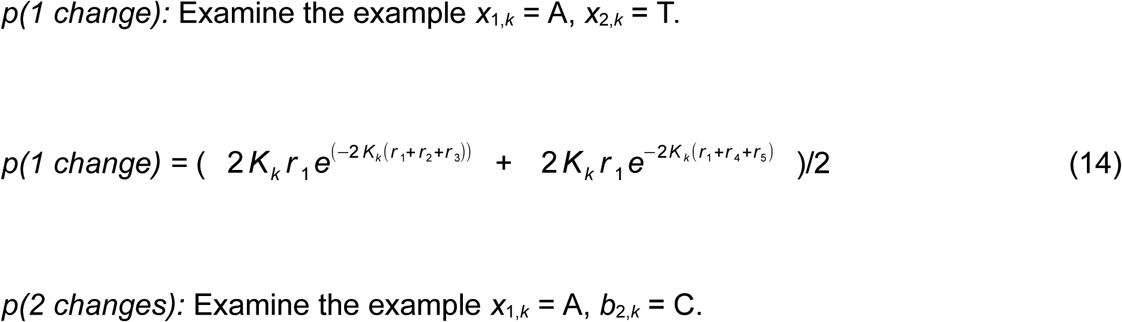

Assuming the configuration of relative rates shown in Fig. 3b (i.e., only *r*_1_ and *r*_4_ are non-zero) and that A is the ancestral base and C is the evolved base. The sequence of events is therefore an A → T change followed by a T → C change. The probability of this event sequence is obtained from:

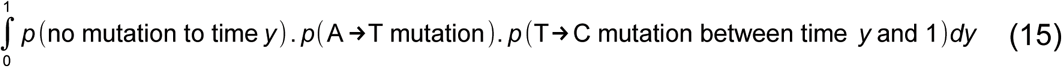

For the example in Fig. 3b, this is:

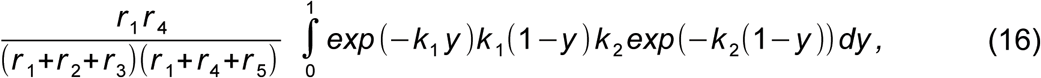

where *k*_1_ and *k*_2_ have the same meanings as above.

The algorithm can be extended to cases where the relative rates are all non-zero.

### Computing uSFS elements

The ML approach described by Keightley et al. (2016) estimates the proportion of density, *π*_*j*_, attributable to the major allele being the ancestral allele *versus* the major allele being the derived allele for each uSFS element pair (indexed by *j* and *n - j*, where *n* is the number of gene copies sampled). We implemented this algorithm as follows, conditional on the ML estimate of the rate parameters, 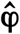(obtained by evaluating equation 1), which are therefore assumed to be known without error. For a uSFS containing *n* elements, *n/2* ML estimates require to be made. Assuming sites evolve independently (cf. equation 1), the likelihood of *π*_*j*_ for the subset of sites (numbering *sitesj)* having *j* copies of the minor allele in the focal species is:

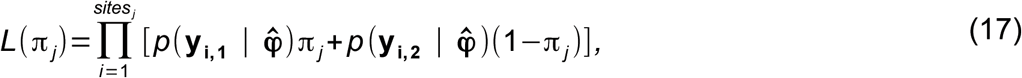

where the probability of the observed nucleotide configuration for the focal species and the outgroups at the site is given by equation (2), evaluated with the major allele 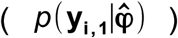 and the minor allele 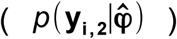 as the state of the focal species at that site (see Fig. 2).

### Computing ancestral state probabilities on a site-by-site basis

The probability of allele X_*i*_ *versus* allele Y_*i*_ being ancestral at site *i* could be computed from their relative probabilities, i.e., 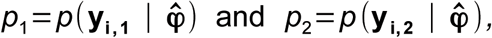 but this only uses information from the estimated rate parameters. It does not incorporate information from the number of major *versus* minor copies at the site. For example, if the outgroup information were uninformative (as in Fig. 1), we would assign *p*_1_ = *p*_2_. If there are few sites in the data set as a whole where the derived allele is at a high frequency, however, the estimated uSFS would tell us that A is more likely to be ancestral.

To infer the ancestral state probabilities for site *i*, information from the estimated rate parameters is augmented by the nearly independent information from the estimated uSFS (cf. Halligan et al. 2013). If there are *j* copies of the minor allele in the focal species at a site *i*, the probability of the major allele X_*i*_ being ancestral is:

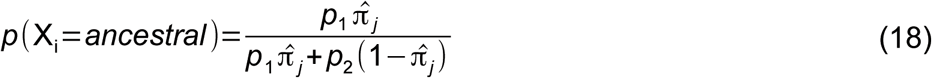

As a check on this equation, it can be shown that the sums of the ancestral state probabilities recovers the estimated uSFS.

### Simulations

We extended a simulation program described by Keightley et al. (2016). This simulates three outgroups for the topology illustrated in Fig. 2. Briefly, unlinked sites were simulated with four nucleotide states in a diploid population of size *N* = 100. The mutation rate per site per generation was set to *μ* = *θ/N*, and the neutral genetic diversity, *θ*, typically set to 0.01. The simulations allowed any variation within a population at a node of the phylogenetic tree (Fig. 2) to be passed to two ancestral sub-populations, which were formed by sampling chromosomes with replacement in one generation. We either simulated neutral sites, or a mixture of neutral and selectively constrained sites. If a mutation occurred at a selectively constrained site, its selection coefficient was *s*/2, where *s* is the difference in fitness between the homozygous mutant and the heterozygote. Fitness effects were multiplicative between and within loci.

### 1000 Genomes data

We downloaded variant calls from the phase3 release of the 1000 Genomes Project from ftp://ftp.1000genomes.ebi.ac.uk/vol1/ftp/release/20130502/ and extracted the 99 unrelated individuals from the Luhya in Webuye, Kenya (henceforth LWK) population. First, we restricted our analyses to sites that were fourfold degenerate in all transcripts of protein-coding genes in humans according to Ensembl release 71. We used the 6-way EPO multiple alignments of primate species, available from ftp://ftp.ensembl.org/pub/release-71/emf/ensembl-compara/epo6primate/ to determine the alleles in orangutans and macaques at each fourfold degenerate site, and to determine whether those sites were within a CpG in humans or either of the outgroup species. The EPO multiple alignments were first converted from .emf format to .maf format, and then specific regions were accessed using the WGAbed package (https://github.com/henryjuho/WGAbed). The data for the human ancestral alleles, as used by the 1000 Genomes Project (1000 Genomes Project Consortium, 2015), were downloaded from ftp://ftp.ensembl.org/pub/release-74/fasta/ancestralalleles/.

Sites were retained for analysis if there was no missing data in humans or either outgroup species. Sites were further assigned to CpG and non-CpG categories. CpG sites were defined as sites that were CpG in their context in any of the three species: humans (including both REF and ALT alleles), orangutans or macaques. Non-CpG sites were defined as sites that were never CpG in their context in any of the same species, including both REF and ALT alleles in the human sample. Alleles at polymorphic sites were used to populate the unfolded site frequency spectrum following two methods. 1) Using the ancestral allele provided by the 1000 genomes project to polarize derived and ancestral variants. 2) using the ML method described in the present study.

## Results

### Simulation results

The method for uSFS inference allows several outgroups to be included in the analysis, but the extent of any benefit from including additional outgroups has been unknown. To investigate this, we simulated unlinked sites according to the tree topology shown in Fig. 2 with three outgroups, recorded the “true” uSFS, and compared it to uSFSs estimated using one, two or three outgroups. High derived allele frequency uSFS elements are expected to be the most affected by misinference (Baudry and Depaulis 2003; Keightley et al. 2016), and contribute most strongly to estimates of positive selection, so we focused on the last uSFS element for polymorphic sites (element 19 of a 20 element uSFS). Our measures of bias and accuracy were the average deviations from the true uSFS elements and root mean-squared error (RMSE) for this element, respectively.

For the case of all sites evolving neutrally, the results (Fig. 4) suggest that adding additional outgroups slightly increases the amount of bias (i.e., high frequency uSFS elements tend to be slightly under-estimated; Fig. 4b). This may be a consequence of violation of the assumption that the phylogenetic tree is known with certainty. The increase in bias is accompanied by reduced RMSE (Fig. 4a) if a second outgroup is added, but there is little benefit from adding a third outgroup. As expected, parsimony-based inference leads to a seriously upwardly biased estimates of the frequency of high frequency derived alleles.

**Figure 4.**
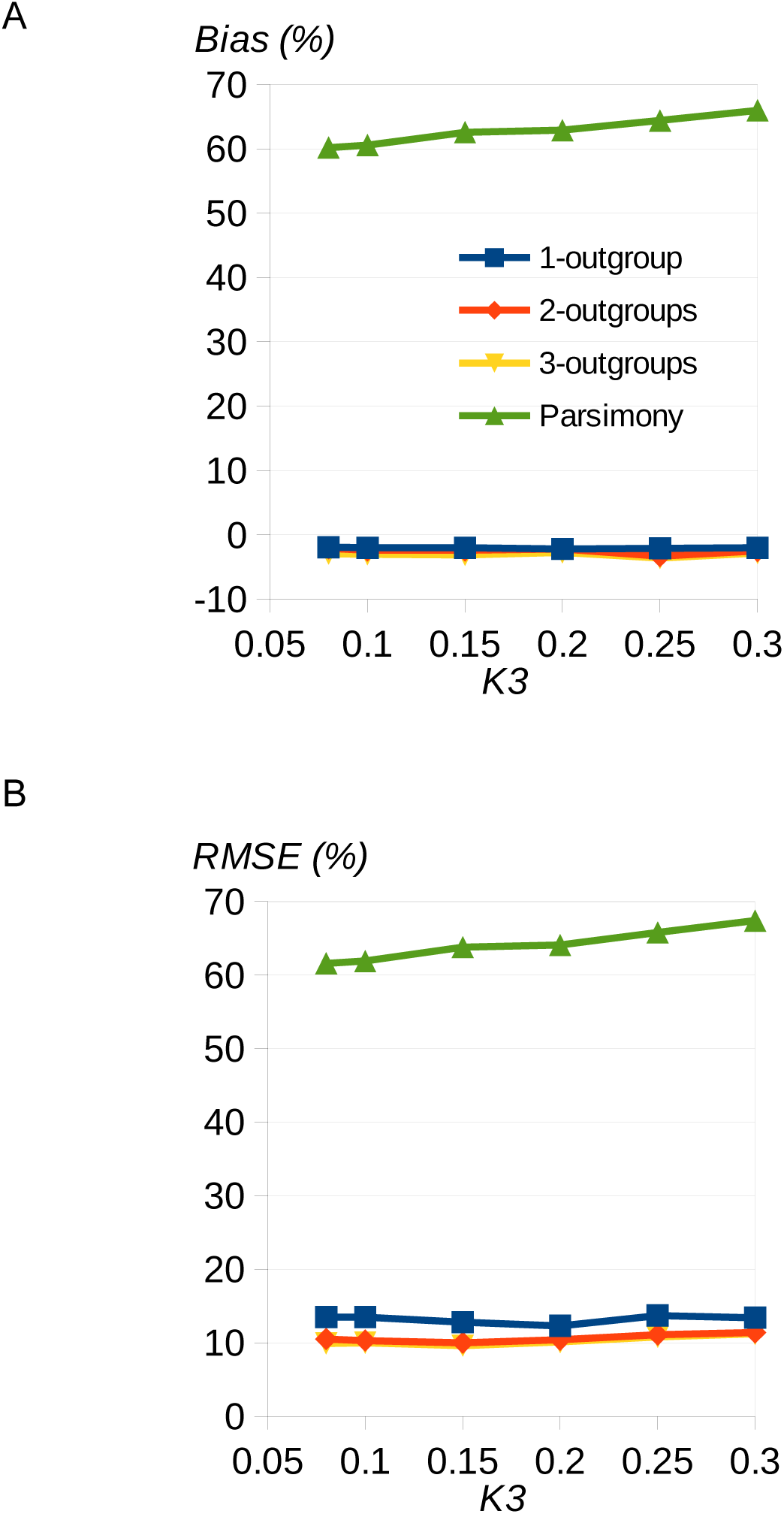
Effect of adding additional outgroups. Simulation results showing A the percentage bias = average deviation from the true uSFS, and B root mean squared error for uSFS element 19, as a function of divergence, *K*_3_, from a third outgroup. There were 100,000 sites simulated in 360 replicates replicate under the JC model, and *K*_1_ = 0.1 and *K*_2_ = 0.1. There were 20 gene copies sampled at each site in the focal species. Blue, red, yellow, green = results from uSFS inference with 1, 2 and 3 outgroups and parsimony, respectively.

We then investigated accuracy and bias for the case of a fraction (*C*) of sites subject to moderately strong purifying selection (scaled selection strength *Ns* = 10). This is relevant for inferring the uSFS for non-neutral sites, such as nonsynonymous sites of protein-coding genes, and for cases where there is variation in the mutation rate among sites leading to variation in the rate of substitution. Such variation violates an assumption of the uSFS inference method, and is therefore expected to cause the method to break down to some extent. As we previously observed (Keightley et al. 2016), the presence of variation in the rate of substitution leads to over-estimation of high derived allele frequency uSFS elements (Fig. 5a). The bias can be serious if there is only one outgroup, but is substantially reduced if a second outgroup is included. However, there is only a small additional benefit from adding a third outgroup. Variation about the observed values is lower, on average, if additional outgroups are included (i.e., RMSE is lower; Fig. 5b), but again adding a third outgroup is of little benefit compared to having just two. As expected, parsimony performs extremely poorly, overestimating the high frequency derived allele frequency.

**Figure 5.**
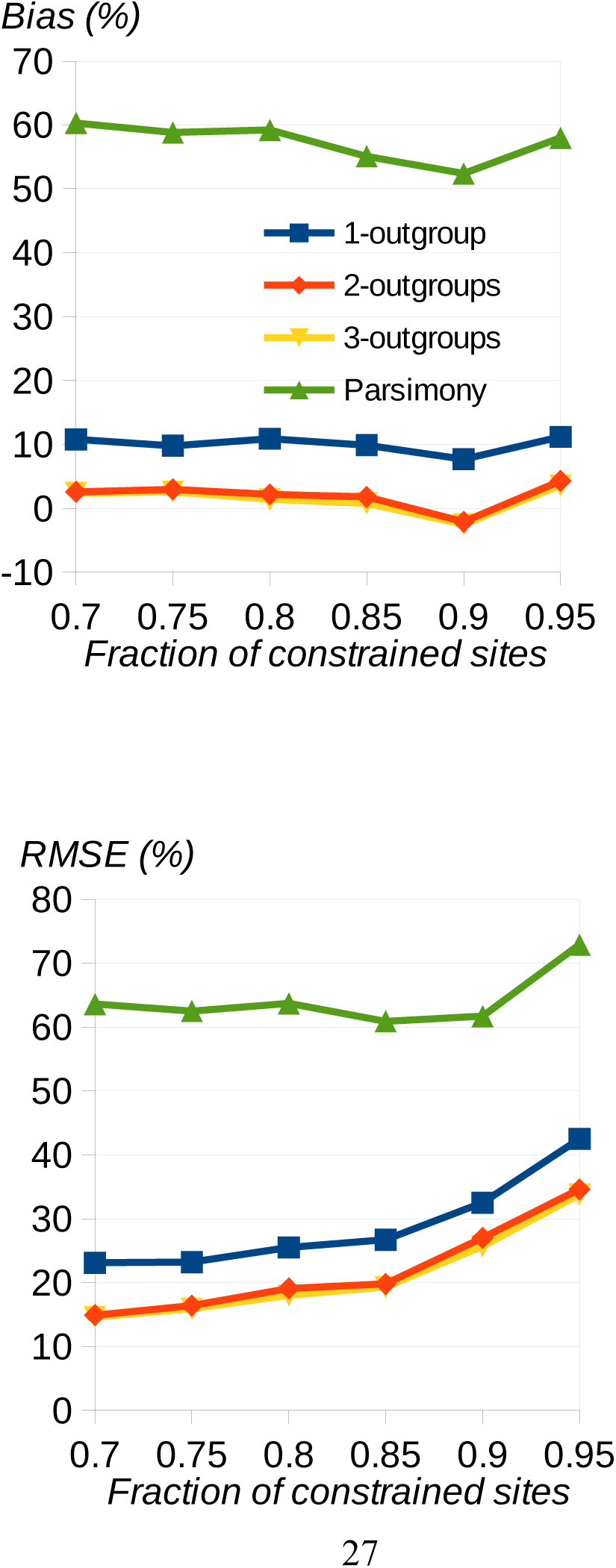
Effect of presence of selectively constrained sites on uSFS inference. Simulation results showing (A) the percentage bias and (B) root mean squared error for uSFS element 19 as a function of the fraction of constrained sites. There were 10,000 sites simulated in 3,600 replicates under the JC model with three outgroups, and *K*_1_ = 0.1, *K*_2_ = 0.15 and *K*_3_ = 0.15. There were 20 gene copies sampled at each site in the focal species. Blue, red, yellow, green = results from uSFS inference with 1, 2 and 3 outgroups and parsimony, respectively.

### Analysis of DPGP phase 2 data

To assess the performance of the uSFS inference procedure in a more realistic situation, we analysed 4-fold-degenerate sites from the Rwandan sequences of the DPGP phase 2, which comprises 17 haploid genomes (provided by J. Campos). We compared the inferred uSFSs obtained using *D. simulans* as the sole outgroup and using both *D. simulans* and *D. yakuba* as outgroups, and investigated the consequences of increasing the complexity of the substitution model. More complex substitution models fit the data much better (Table 1), largely driven by the approximately two-fold transition:transversion mutation bias, captured by the K2 model.

**Table 1.** Differences between log likelihood of simpler models and the R6 model. The data analyzed are 4-fold-degenerate sites from the Rwandan sequences of DPGP phase 2, using *D. simulans* and *D. yakuba* as outgroups.

| *Model* | *ΔLog likelihood* |
| --- | --- |
| Jukes-Cantor | -13,000 |
| Kimura 2-parameter | -1,400 |
| R6 | 0 |

Although different nucleotide substitution models produce large differences in log likelihood, the estimated uSFS is appreciably different only between JC and K2 model, and indistinguishable between the K2 and R6 models (Fig. 6a). Consistent with the simulation results, the inclusion of a second outgroup (*D. yakuba*) perceptibly reduced the high derived allele frequency uSFS elements, compared to using a single outgroup (*D. simulans*) (Fig. 6b). There is an uptick at the right hand side of the inferred uSFS, but it is unknown whether this is a consequence of mis-inference or ongoing positive selection on 4-fold sites or positive selection on linked sites. Consistent with the simulations, parsimony infers a substantially higher frequency of high frequency derived allele classes.

**Figure 6.**
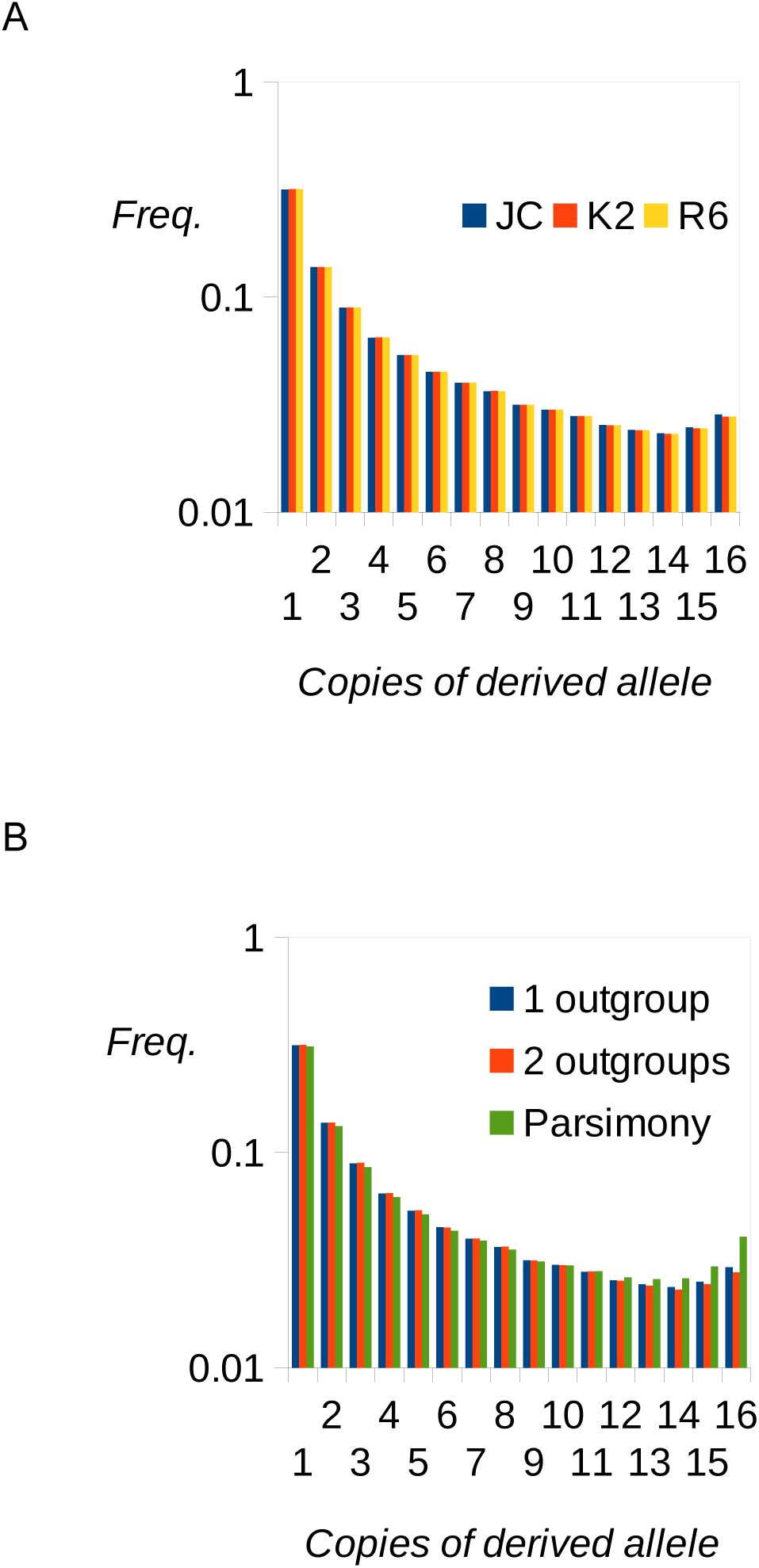
Analysis of 4-fold-degenerate sites of DPGP phase 2. (A) uSFSs estimated assuming three different substitution models. (B) uSFSs estimated using the method described in this paper based on one outgroup (*D. simulans*) or two outgroups (*D. simulans* and *D. yakuba*) along with the uSFS inferred using parsimony.

### Analysis of 1000 Genomes data

SNP ancestral states inferred by the 1000 Genomes Project Consortium (2010; 2015) have been widely used (e.g., Mondal et al. 2015; Yang, and Slatkin 2016; Harris and Pritchard 2017). In their 2015 paper, a heuristic approach was used to assign the ancestral state based on the inferred human-chimp common ancestor and the human-chimp-orangutan common ancestor. Allele frequency information was not incorporated. We re-inferred the ancestral state at 4-fold-degenerate sites in the LWK population using the ML method presented here, and compared the resulting uSFSs (Fig. 7). Because uSFSs from the full dataset of 99 individuals (198 chromosomes) were difficult to visualise, we downsampled the LWK population to 25 randomly chosen individuals. The results from the full dataset are qualitatively similar to those from the downsampled data and are presented in Fig S1.

**Figure 7.**
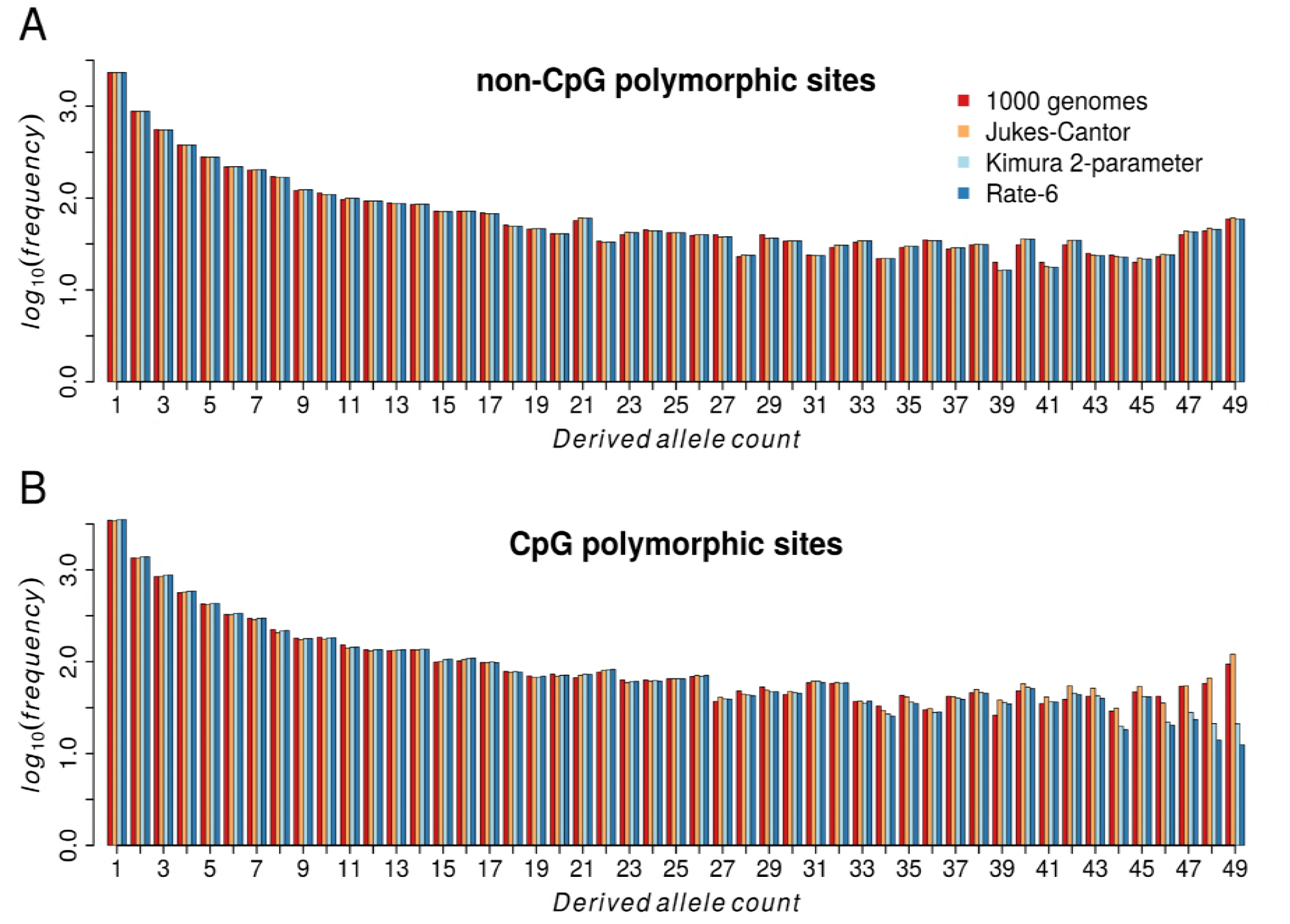
uSFSs inferred by the 1000 Genomes Project and by the methods described in this paper for three nucleotide substitution models. (A) non-CpG sites. (B) CpG sites.

For non-CpG sites the uSFS produced using the 1000 Genomes Project’s ancestral states and the uSFSs produced by our ML method broadly agreed (Fig. 7a). In contrast, for CpG sites, the results under the 1000 genomes method and the Jukes-Cantor substitution model depart from the Kimura 2-parameter model and the R6 model at the right hand side of the inferred uSFS (Fig. 7b). Under 1000 Genomes and JC, there is a pronounced uptick at high frequency derived variants which is not present in the two more complex substitution models.

CpG sites have a ~10-fold higher mutation rate than non CpG sites in humans, due to an elevation in the number of C → T and G → A transitions (Nachman and Crowell, 2000). This was borne out in the inferred branch lengths and the ratio of transition rate to transversion rate (*κ*) at the two classes of site. Under the R6 model, which is the best-fitting model for both classes of site, the length of the branch between the Human-Orangutan common ancestor and Humans was 0.0083 for non-CpG sites and 0.092 for CpG sites. Estimates of *κ* under the Kimura 2-paramter model were 4.2 and 8.3 for non-CpG and CpG sites, respectively, which are broadly in agreement with previous studies (e.g., Keightley et al. 2011).

## Discussion

In this paper, we have generalized a method we developed for inferring the uSFS (Keightley et al. 2016) to allow the inclusion of multiple outgroup species and potentially any phylogenetic tree topology (although only topologies of the type illustrated in Fig. 2 have been implemented in the software). We have also implemented three substitution models: the Jukes-Cantor (JC) and Kimura 2-parameter (K2) models, and the “R6” model with six symmetrical relative mutation rates. These models are nested. The K2 model gives the same likelihood as the JC model if the transition:transversion ratio parameter *κ* is fixed at 1. If the R6 parameters are constrained such that *r*_3_ = *r*_4_ (transition mutations) and *r*_1_ = *r*_2_ = *r*_5_ = *r*_6_ (transversion mutations) (see Fig. 3), the same maximum likelihood is obtained as the K2 model. Consistent with our previous results (Keightley et al. 2016), simulations suggest that the inclusion of a second outgroup generally increases the accuracy of uSFS inference, especially in the presence of variation in the rate of substitution among sites. The inclusion of a third outgroup did not, however, lead to a further improvement in uSFS inference accuracy. In the real data sets we have analyzed from Drosophila and humans, more complex substitution models gave substantially higher log likelihoods in stage 1 of the analysis (evolutionary rate parameter estimation), but this did not translate into a benefit in stage 2 (uSFS element inference) beyond the K2 model. The nucleotide substitutions models implemented are somewhat simplified in the sense that rates of change between pairs of nucleotides are symmetrical and these parameters do not vary between branches. It is possible that more complex models allowing these complications would lead to a further improvement, given that such effects are common in real data. A further weakness we hope to address in the future is its non-context dependence of substitution model (so we cannot deal with hypermutable CpGs), and further development along the lines of, for example, Arndt et al. (2003) will be needed.

In the present study we divided the data from the 1000 genomes project into CpG and non-CpG sites, and inferred uSFSs separately for each class. At non-CpG sites there was a close agreement between the uSFS generated using the 1000 Genomes Project’s ancestral alleles to polarize variants and the uSFSs generated using the ML method. Parsimony is a more justifiable method of reconstructing ancestral states when the amount of change is small over the evolutionary time being considered, because it assumes *a priori* that change is unlikely (Felsenstein, 1981). In contrast, parsimony is likely to be less accurate at CpG sites, which have a ~10-fold higher rate of evolution. Our results bear this out. The uSFSs for CpG sites differed in the frequency of high frequency derived variants between the 1000 Genomes and the K2 and R6 models. These are the class of variants where the greatest probability of misinference is expected. The JC model more closely mirrored the 1000 Genomes Project uSFS, which may be because it was unable to capture the ratio between transition rate and transversion rate at CpG sites, which is ~two-fold more extreme compared to non-CpG sites.

We have also addressed the problem of calculating ancestral state probabilities for polymorphic sites on a site-by-site basis. In doing so, we take into account both the nucleotide substitution parameter estimates (which determine the frequencies of multiple hits) and the frequencies of derived *versus* ancestral alleles at other sites in the data. There are two main situations where this can make a significant difference compared to using of parsimony. The first concerns sites where the outgroups are different in state from the focal species. These sites are frequently removed from the analysis (e.g., Keinan et al. 2007; Sabeti et al. 2007; Langley et. 2012; 1000 Genomes Project Consortium 2010; 2015), but will lead to an under-representation of polymorphic sites, especially sites that have a low frequency of the derived allele, which tend to be the most common. The second situation concerns to the bias of parsimony, leading to over-estimation of the frequency of sites with a high frequency of the derived allele. Consider the two configurations of nucleotides at a site of focal species and two outgroups shown in Fig. 8. This is one of a large number of sites generated by simulation for which the uSFS has been estimated. At the site in question, there are 19 As and 1 T in the 20 gene copies sampled. In the left-hand panel (Fig. 8a), the two outgroups are state A. By parsimony, the ancestral allele of the variation in the focal species would therefore be assigned as A. If the branch length *b*_1_ (Fig. 2) is 0.05, and using only information from the inferred substitution rates (i.e., using the relative values of *p*_1_ and *p*_2_ calculated using equation 2), *p*(A = ancestral) is 0.98. Taking into account the fact that high frequency derived allele sites are rare in the data set as a whole, and applying equation (18), base A is even more strongly supported as the ancestral allele, i.e., *p*(A = ancestral) >0.99. This illustrates that parsimony is a good approximation for sites likely to have a low number of derived gene copies. The outcome is different for Fig. 8b, where the two outgroups have the same state as the minor allele of the focal species. By parsimony, the ancestral allele would be assigned C, implying that we are certain the site has 19 copies of the derived allele. Using only information from substitution rate parameters and applying equation (2), *p*(A = ancestral) is 0.016, which is close to the result using parsimony. Taking into account other sites in the data, which tell us that sites having 19 derived allele copies are uncommon, and applying equation (18), *p*(A = ancestral) is 0.14. Thus, we are a lot less certain that the derived allele is A at this site. This probability rises (drops) if the outgroups are more distant (closer) to the focal species.

**Figure 8.**
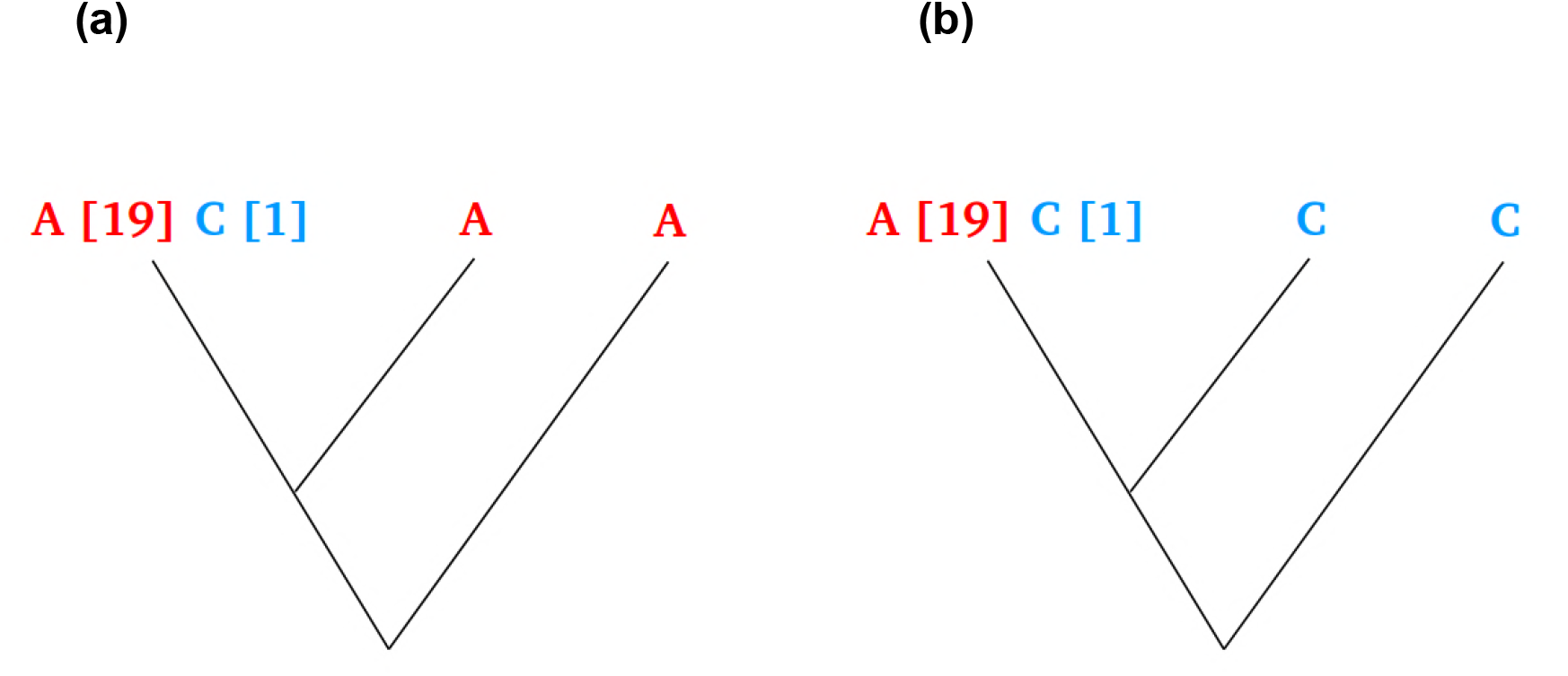
Example of a polymorphic site where 20 gene copies are sampled in a focal species and two outgroups have different nucleotide states.

## Acknowledgements

This project has received funding from the European Research Council (ERC) under the European Union’s Horizon 2020 research and innovation programme (grant agreement No. 694212). We thank Tom Booker and Dan Halligan for helpful discussions.

## Software availability

A program to carry out ML uSFS inference and computation of ancestral state probabilities is available from PDK’s website.

## References

Arndt, P. F., Burge, C. B. and Hwa, T. 2003 DNA sequence evolution with neighbor-dependent mutation. Journal of Computational Biology 10: 313–322.

Baudry E, Depaulis, F. 2003 Effect of misoriented sites on neutrality tests with outgroup. Genetics 165: 1619–1622.

Boyko, A. R., Williamson, S. H., Indap, A. R., Degenhardt, J. D., Hernandez, R. D., et al. 2008 Assessing the evolutionary impact of amino acid mutations in the human genome. PLoS Genet. 4: e1000083.

Collins, TM, Wimberger, PH, Naylor, GJP 1994 Compositional bias, character state bias and character state reconstruction using parsimony. Syst. Biol. 43: 482–496

Dreszer, T. R., Wall, G. D., Haussler, D. and Pollard, K. S. 2007 Biased clustered substitutions in the human genome: The footprints of male-driven biased gene conversion. Genome Res. 17: 1420–1430.

Eyre-Walker, A. 1998 Problems with parsimony in sequences of biased base composition. J. Mol. Evol. 47: 686–690.

Fay, J. C. and Wu, C.-I. 2000 Hitchhiking under positive Darwinian selection. Genetics 155: 1405–1413.

Felsenstein, J. 1981 Evolutionary trees from DNA sequences: a maximum likelihood approach. J. Mol. Evol. 17: 368–376.

Halligan, D. L., Kousathanas, A., Ness, R. W., Harr, B., Eory, L., Keane, T. M., Adams, D. J. and Keightley, P. D. (2013). Contributions of protein-coding and regulatory change to adaptive molecular evolution in murid rodents. PLoS Genet. 193: 1197–1208.

Harris, K. and J. K Pritchard 2017 Rapid evolution of the human mutation spectrum. eLife 6: e24284.

Hernandez, RD, Williamson, SH and Bustamante, CD 2007 Context dependence, ancestral misidentification, and spurious signatures of natural selection. Mol. Biol. Evol. 24: 1792–1800.

Jackson, B. C., Campos, J. L., Haddrill, P. R., Charlesworth, B. and Zeng, K. 2017 Variation in the intensity of selection on codon bias over time causes contrasting patterns of base composition evolution in *Drosophila*. Genome Biol. Evol. 9: 102–123.

Keightley, P. D., Eöry, L., Halligan, D. L. and Kirkpatrick, M. (2011). Inference of mutation parameters and selective constraint in mammalian coding sequences by approximate Bayesian computation. Genetics 187: 1153–1161.

Keightley, P. D., Campos, J. L., Booker T. R. and Charlesworth, B. 2016 Inferring the frequency spectrum of derived variants to quantify adaptive molecular evolution in protein-coding genes of *Drosophila melanogaster*. Genetics 203: 975–984.

Keinan, A., Mullikin, J. C., Patterson, N. and Reich, D. 2007 Measurement of the human allele frequency spectrum demonstrates greater genetic drift in East Asians than in Europeans. Nature Genet. 39: 1251–1255.

Langley, C. H., Stevens, K., Cardeno, C., Lee, Y. C. G., Schrider, D. R., et al. 2012 Genomic Variation in Natural Populations of *Drosophila melanogaster*. Genetics 192: 533–598.

Lohse, R. J. Harrison and N. H. Barton 2011 A general method for calculating likelihoods under the coalescent process. Genetics 189: 977–987.

Matsumoto, T., Akashi, H. and Yang, Z. 2015 Evaluation of ancestral sequence reconstruction methods to infer nonstationary patterns of nucleotide substitution. Genetics 200: 873–890.

Mondal, M., Casals, F., Xu, T., Dall’Olio, G. M., Pybus, M. et al. 2016 Genomic analysis of Andamanese provides insights into ancient human migration into Asia and adaptation. Nature Genet. 48: 1066–1070.

Nachman, M. W. and Crowell, S. L. 2000 Estimate of the mutation rate per nucleotide in humans. Genetics 156: 297–304.

Nelder, J. A. and Mead, R. 1965 A simplex method for function minimization. Comput. J. 7: 308–313.

Sabeti, P. C., Varilly, P., Fry, B., Lohmueller, J., Hostetter, E. et al. 2007. Genome-wide detection and characterization of positive selection in human populations. Nature 449: 913–918.

Schmidt, J. M., Battlay, P., Gledhill-Smith, R. S., Good, R. T. Lumb, C., Fournier-Level, A. and Robin, C. 2017. Insights into DDT Resistance from the Drosophila melanogaster Genetic Reference Panel. Genetics 207: 1181–1193.

Schneider, A., Charlesworth, B., Eyre-Walker, A. and Keightley, P. D. 2011 A method for inferring the rate of occurrence and fitness effects of advantageous mutations. Genetics 189: 1427–1437.

Tataru, P., Mollion, M., Glémin, S. and Bataillon, T. 2017 Inference of distribution of fitness effects and proportion of adaptive substitutions from polymorphism data. Genetics Early online September 25, 2017; https://doi.org/10.1534/genetics.117.300323

The 1000 Genomes Project Consortium 2010 A map of human genome variation from population-scale sequencing. Nature 467: 1061–1073.

The 1000 Genomes Project Consortium 2015 A global reference for human genetic variation. Nature 526: 68–74.

Voight, BF, Kudaravalli, S, Wen, X, Pritchard, JK 2006 A map of recent positive selection in the human genome. PLoS Biol. 4: e72.

Yang, M. A. and M. Slatkin 2016 Using ancient samples in projection analysis. G3: Genes, Genomes, Genetics 6: 99–105.

Yang, Z. 2007 PAML 4: Phylogenetic Analysis by Maximum Likelihood. Mol. Biol. Evol. 24: 1586–1591.

Zeng, K., Y.-X. Fu, S. Shi, and C.-I. Wu. 2006 Statistical tests for detecting positive selection by utilizing high-frequency variants. Genetics 174:1431–1439, 2006.

